# A genome-scale CRISPR interference guide library enables comprehensive phenotypic profiling in yeast

**DOI:** 10.1101/2020.03.11.988105

**Authors:** Nicholas J. McGlincy, Zuriah A. Meacham, Kendra Swain, Ryan Muller, Rachel Baum, Nicholas T. Ingolia

## Abstract

CRISPR/Cas9-mediated transcriptional interference (CRISPRi) enables programmable gene knock-down, yielding interpretable loss-of-function phenotypes for nearly any gene. Effective, inducible CRISPRi has been demonstrated in budding yeast, but no genome-scale guide libraries have been reported. We present a comprehensive yeast CRISPRi library, based on empirical design rules, containing 10 distinct guides for most genes. Competitive growth after pooled transformation revealed strong fitness defects for most essential genes, verifying that the library provides comprehensive genome coverage. We used the relative growth defects caused by different guides targeting essential genes to further refine yeast CRISPRi design rules. In order to obtain more accurate and robust guide abundance measurements in pooled screens, we link guides with random nucleotide barcodes and carry out linear amplification by in vitro transcription. Taken together, we demonstrate a broadly useful platform for comprehensive, high-precision CRISPRi screening in yeast.

## INTRODUCTION

Systematic genetic analysis — the comprehensive assessment of phenotypes across a large and defined collection of genetic perturbations — is a powerful approach for learning the organizing principles of molecular and cellular processes. Systematic analyses provide quantitative phenotypic profiles that serve as a rich and nuanced source of information, as well as identifying key candidate genes in the manner of a classical genetic screen. Truly comprehensive, systematic analysis was realized first in budding yeast (*Saccharomyces cerevisiae*), with the creation of the deletion collection, an arrayed library of ~6,000 yeast strains that each contain one barcoded gene knock-out (Giaever et al, 2002; Winzeler et al, 1999). Subsequently, RNA interference was harnessed for large-scale genetic analysis in many organisms (Fraser et al, 2000; Gonczy et al, 2000) and cell models (Paddison et al, 2004). More recently, programmable RNA-guided DNA targeting by Cas9 and other CRISPR-associated proteins has emerged as an enabling technology for systematic genetic analysis. In its native form, Cas9 cleaves DNA at sites complementary to a short guide RNA (Jiang & Doudna, 2017), often leading to mutations mediated by error-prone repair pathways (Jasin & Haber, 2016). Guide RNA libraries thereby enable comprehensive, targeted mutagenesis that offers advantages for comprehensive genetic screening (Morgens et al, 2016).

Catalytically inactive Cas9 (dCas9) retains RNA-guided DNA binding activity that can be harnessed for many other purposes. When dCas9 is fused with another protein, it targets this fusion partner to the genomic sequence specified by a guide RNA, enabling an array of novel approaches to measure and manipulate the genome (Dominguez et al, 2016). Targeting co-repressor proteins to eukaryotic promoters leads to CRISPR-mediated transcriptional interference (CRISPRi), a powerful and general approach to reduce transcription from the targeted locus (Gilbert et al, 2013). CRISPRi yields reproducible, partial loss-of-function phenotypes that offer advantages over Cas9-mediated cutting in systematic genetic analysis (Gilbert et al, 2014; Kampmann, 2018). Essential genes can be analyzed easily with CRISPRi, and knock-down can be quickly activated and quickly relieved by conditional expression of the dCas9 fusion protein or the guide RNA. Genome-wide CRISPRi libraries are thus highly desirable even in budding yeast, where deletion collections and other resources are available.

Although no comprehensive guide libraries have been reported for budding yeast, we do have optimized tools to support CRISPRi screening. Transcriptional interference by dCas9-mediated recruitment of repressor domains was pioneered in yeast, and potent CRISPRi has been achieved with a dCas9-Mxi1 fusion that links dCas9 with a fragment of a mammalian repressor (Gilbert et al., 2013). Single guide RNAs (sgRNAs) can be expressed from an RNA Polymerase III promoter taken from the yeast RPR1 gene. Furthermore, embedding tetracycline operator (tetO) sites in this promoter confers tetracycline-inducible guide expression, and thus regulated CRISPRi activity (Farzadfard et al, 2013). This inducible guide expression system has been used to create substantial collections of effective guides spanning almost 500 genes (Jaffe et al, 2019; Smith et al, 2016), which have provided rules for guide RNA design (Smith et al., 2016). In yeast, as in many eukaryotes, chromatin accessibility at a target DNA sequence and the position of this sequence relative to the transcription start site are key determinants of effective CRISPRi. Guides binding nucleosome-free sites in the region 200 bp just upstream of the transcription start site were most likely to be active; although these two factors are correlated, each appears to be important individually.

Using these rules, we have generated and validated a genome-wide CRISPRi screening system for budding yeast. We first constructed a comprehensive library of episomal guide expression plasmids. In order to quantify guide abundance in screens, we link guide RNAs with random nucleotide barcodes and amplify these barcodes by in vitro transcription. We used this barcoded guide library to carry out a pooled growth screen in a continuous culture of prototrophic yeast in minimal synthetic media. Guides produced distinctive, reproducible fitness effects that could be inferred from exponential dynamics of their abundance during competitive growth. We found guides having strong growth defects for the great majority of essential genes, showing that our library provides excellent coverage. Comparisons of the active and inactive guides allowed us to further refine design rules for yeast CRISPRi and better assign target genes to guide sites at closely-spaced, divergent promoters, which are common in yeast. Our system for high coverage, high efficacy CRISPRi screening provides a broadly useful tool for the budding yeast community with numerous applications.

## RESULTS

### Design of a guide RNA library for genome-wide CRISPRi in budding yeast

We set out to design a library of yeast guide RNAs suitable for genome-wide CRISPRi screening. In yeast, the efficiency of transcriptional interference is affected by the distance between the target sequence and the transcription start site and by the accessibility of the DNA at that target (Smith et al., 2016). Even after controlling for these parameters, only a fraction of guide RNAs inhibit transcription effectively, and so we aimed to select ten guides for each of the annotated genes in the yeast genome (Cherry et al, 2012).

We implemented a deterministic target site selection scheme based on heuristics that seemed likely to pick active and specific guides (Fig. 1a). We prioritized guides first by preferring targets that were unique in the genome and specific to one target gene, and then according to accessibility as determined by ATAC-Seq (Schep et al, 2015). We also ensured that our guides were distributed across the full range of positions where CRISPRi appears effective by selecting at least one target site from each of a few different zones within the overall promoter region (Fig. 1b). When the transcription start site was known from transcript isoform sequencing (Pelechano et al, 2013), we picked targets in a range from 220 base pairs upstream of this transcriptional start through 20 nucleotides downstream. When no transcriptional start site was available, we picked targets between 350 and 30 nucleotides upstream of the coding sequence. Using these rules, we designed 61,094 guides targeting all annotated protein-coding genes, excepting those predicted open reading frames characterized as “dubious”, and also against non-coding RNAs. The majority of genes were targeted by ten unique and unambiguous guides (Fig. 1c).

**Figure 1.**
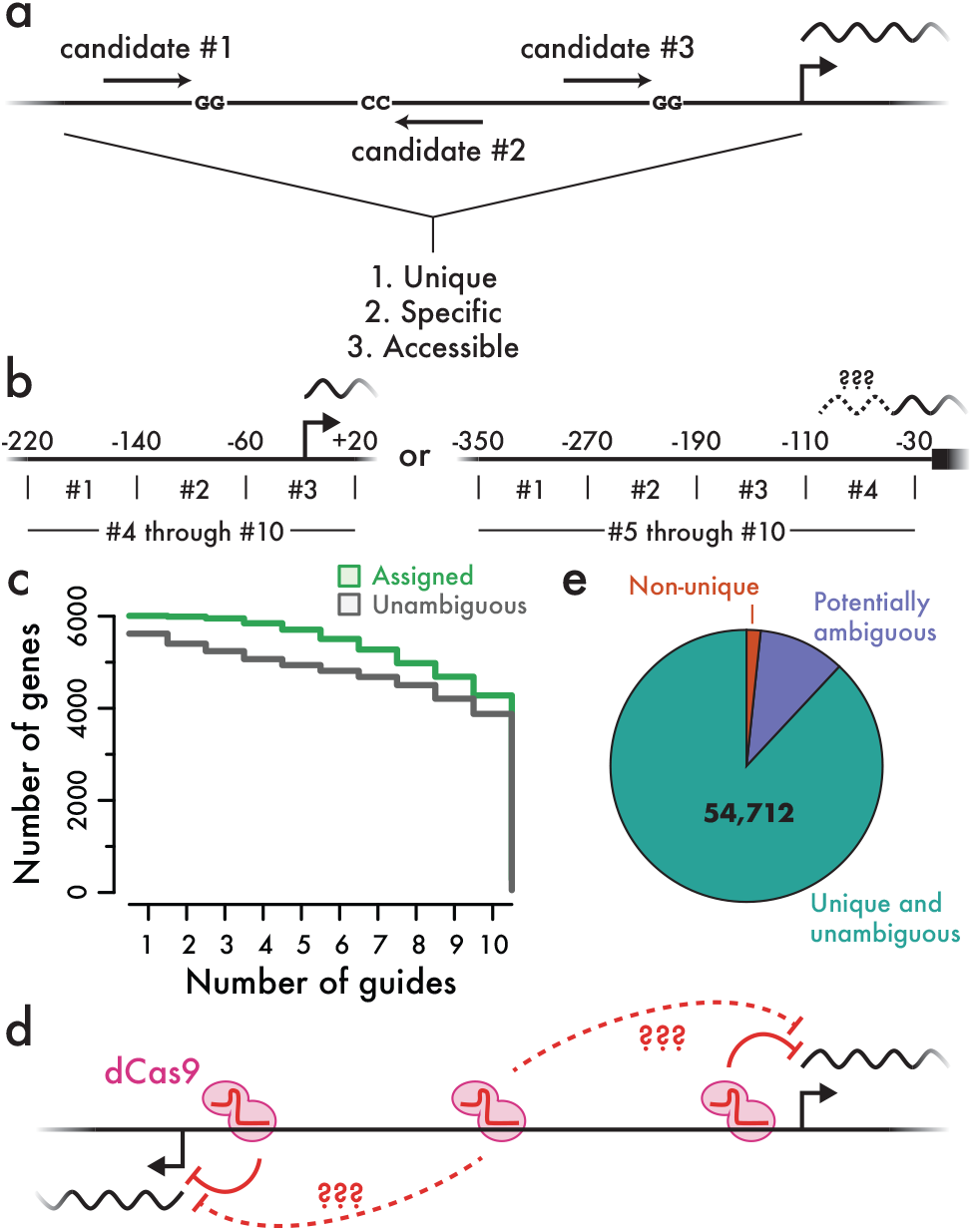
Design of a genome-wide yeast CRISPRi library. **(a)** Candidate guide RNA sites in promoter regions were identified and scored to prioritize target sites that were unique in the genome, specific to one promoter, and located in accessible chromatin. **(b)** Promoter regions targeted by guide RNAs. When transcription start sites were known, we selected one guide each from three separate regions around the transcription start site, along with seven others in the overall promoter. When transcription start sites were not known, we targeted a wider area upstream of the start of the coding sequence. **(c)** Cumulative distribution of genes targeted by up to ten distinct guides. Unambiguous guides do not fall within the potentially active region for any other gene. Assigned guides have a single likely target based on empirical measurements of guide activity and location. **(d)** Schematic of an ambiguous guide at a divergent promoter. **(e)** Classification of designed guides according to unique genomic sites and unambiguous target genes.

The compact yeast genome, containing many divergently transcribed genes separated by only a few hundred base pairs, poses challenges for guide design. Guides falling in the overlapping region between two divergently transcribed promoters have at least the potential to target either gene (Fig. 1d). Roughly 10% of the guides we selected were potentially ambiguous in this way, in addition to a very small fraction of non-unique guide sequences (Fig. 1e). Since the distance between a guide and a promoter is a key determinant of its efficacy, we reasoned that we could assign many of these potentially ambiguous guides to one likely target. As described below, we were able to carry out this assignment using large-scale empirical measures of guide activity, further enhancing our coverage of the genome.

### Linear amplification of guide-linked nucleotide barcodes by in vitro transcription enables precise measurements of guide frequency

Pooled CRISPR screening relies on measurements of guide RNA abundance in a population of cells, typically carried out by high-throughput sequencing (Kampmann, 2018). Phenotypic effects manifest as changes in these guide frequencies caused by competitive growth under different conditions or by flow cytometric sorting for specific phenotypes. We therefore sought the most precise and robust approach to measure the abundance of guide RNA expression plasmids from yeast. Rather than sequencing guides directly, we used arbitrary nucleotide barcodes embedded in the guide RNA expression plasmid. One major advantage of sequencing these barcodes is that each guide can be linked to a few different barcodes, providing replicate measurements of its effect within a single experiment (Michlits et al, 2017; Schmierer et al, 2017). In contrast, direct guide sequencing cannot distinguish between independently transformed lineages within a single experiment. Barcode sequencing also allows us to distinguish defective guide RNA expression constructs, which typically cause no phenotypic effects, from sequencing errors arising during quantitation. We can detect and correct single-nucleotide sequencing errors that we observe when quantifying barcodes while excluding barcodes linked to guides with errors introduced during synthesis or cloning.

High-throughput sequencing of barcodes (or guide RNAs) requires substantial, selective amplification of DNA recovered from cells. Pooled screening approaches typically use populations of 1 million to 100 million cells, each yielding one or a few copies of the DNA to be counted (Kampmann, 2018). High-throughput sequencing requires roughly 10 billion input molecules, and the DNA samples recovered from cell pools are generally amplified at least a thousand-fold to create a sequencing library (van Dijk et al, 2014). Exponential PCR can easily achieve this amplification, but also introduces multiplicative noise, and stochastic events occurring in early PCR cycles are amplified along with the underlying barcode abundances. Linear amplification by in vitro transcription offers an attractive alternative to PCR amplification (Van Gelder et al, 1990) and has been used productively in single-cell DNA and RNA sequencing approaches (Chen et al, 2017; Eberwine et al, 1992; Hashimshony et al, 2012). We confirmed that in vitro transcription of template plasmid isolated from budding yeast yielded ~5,000-fold amplification over a wide range of template DNA amounts and tolerated substantial non-template DNA.

We devised a strategy for measuring barcode abundance by sequencing, using initial linear amplification by in vitro transcription, that substantially reduced noise relative to direct PCR amplification (Fig. 2). The RNA product of in vitro transcription is reverse transcribed back into DNA (IVT-RT), which serves as a template for limited PCR that generates double-stranded DNA with flanking sequences required for high-throughput sequencing (Fig. 2a). In order to validate our IVT-RT library generation strategy and compare it directly with PCR amplification, we transformed yeast with a plasmid library containing ~250,000 random nucleotide barcodes, carried out batch selection for transformants, and recovered plasmid DNA from two replicate samples drawn from this transformed population. IVT-RT libraries generated from these replicate DNA samples showed substantially better quantitative agreement than matched libraries constructed by direct PCR amplification (Fig. 2c, 2d). Duplicate IVT-RT libraries from the same population showed a correlation *r* = 0.98, whereas PCR libraries correlated substantially worse, *r* = 0.93. Dispersion estimates from replicate IVT-RT libraries showed markedly lower variances than matched PCR libraries at equivalent read depth (Fig. 2e), which translates into more precise guide abundance measurements and thus greater statistical power to resolve phenotypic differences.

**Figure 2.**
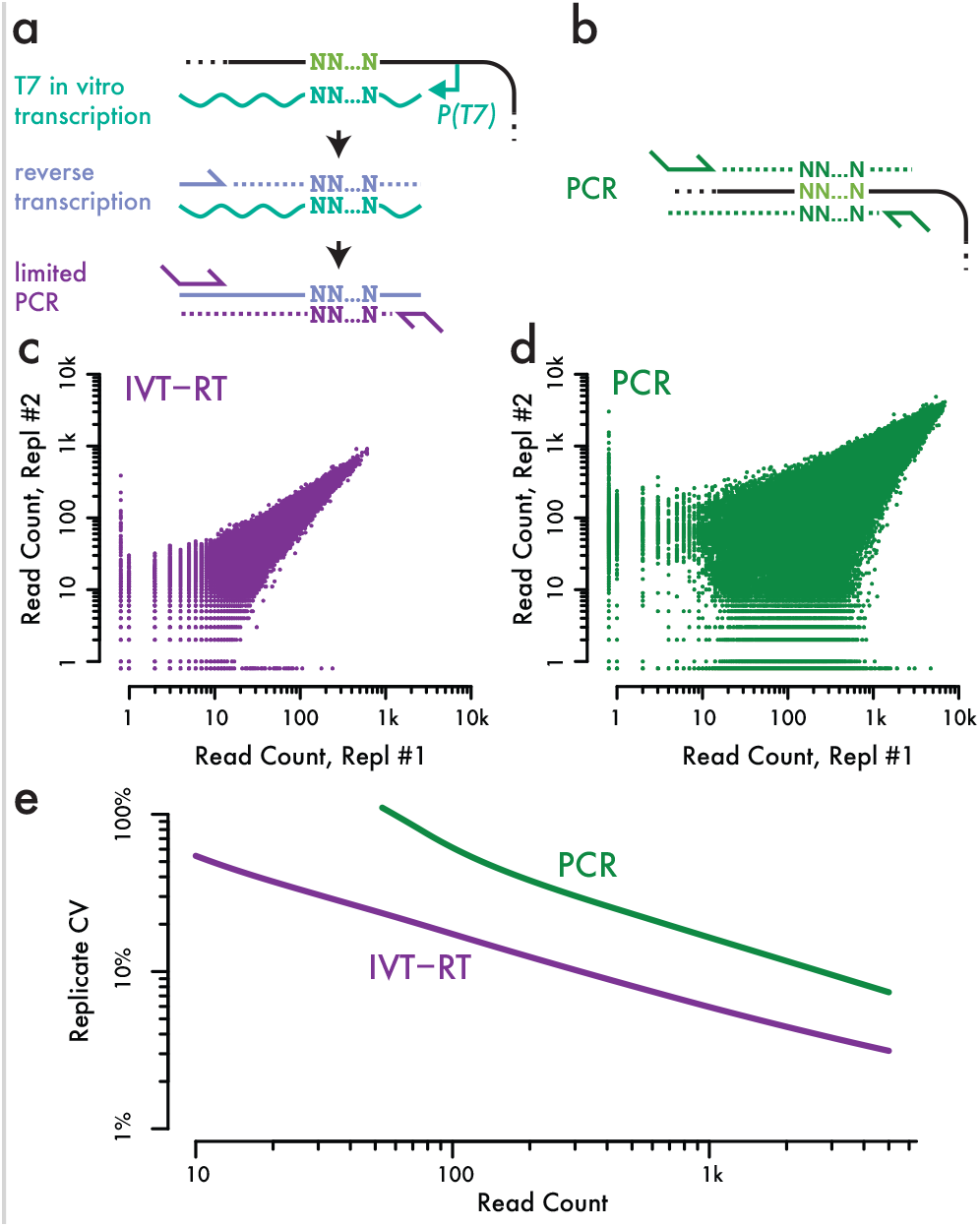
Linear amplification by in vitro transcription improves precision of barcode abundance measurements. **(a, b)** Schematics of barcode library generation by (a) in vitro transcription followed by RT-PCR and (b) direct PCR amplification. **(c, d)** Barcode read counts in libraries prepared from replicate DNA samples by (c) IVT-RT-PCR and (d) direct PCR. **(e)** Dispersion between replicate measurements as a function of read count.

### Construction of a barcoded, genome-wide library of inducible guide RNAs

Based on these observations, we generated a genome-wide yeast CRISPRi guide expression library with linear IVT-RT amplification of linked nucleotide barcodes (Fig. 3a). Our library includes only the guide RNA cassette, and requires separate expression of the rest of the inducible CRISPRi machinery, as we found that smaller plasmids containing just the guide RNAs improved both the diversity of pooled transformations and the yield of subsequent plasmid recovery. We first introduced guide RNAs into a tetracycline-inducible derivative of a RNA polymerase III promoter in a high-efficiency, bacterial cloning reaction that maximized library diversity (Fig. 3b and Figure S1). We then added barcodes in a second cloning step and controlled the yield of the bacterial transformation in order to capture an average of 4 barcodes per guide RNA (Fig. 3b and Figure S2). While greater barcode diversity is beneficial to a point, limiting the number of barcodes allows us to maintain a substantial number of cells per barcode, which is important for robust barcode counting, and to assign barcodes to guide RNAs reliably.

**Figure 3.**
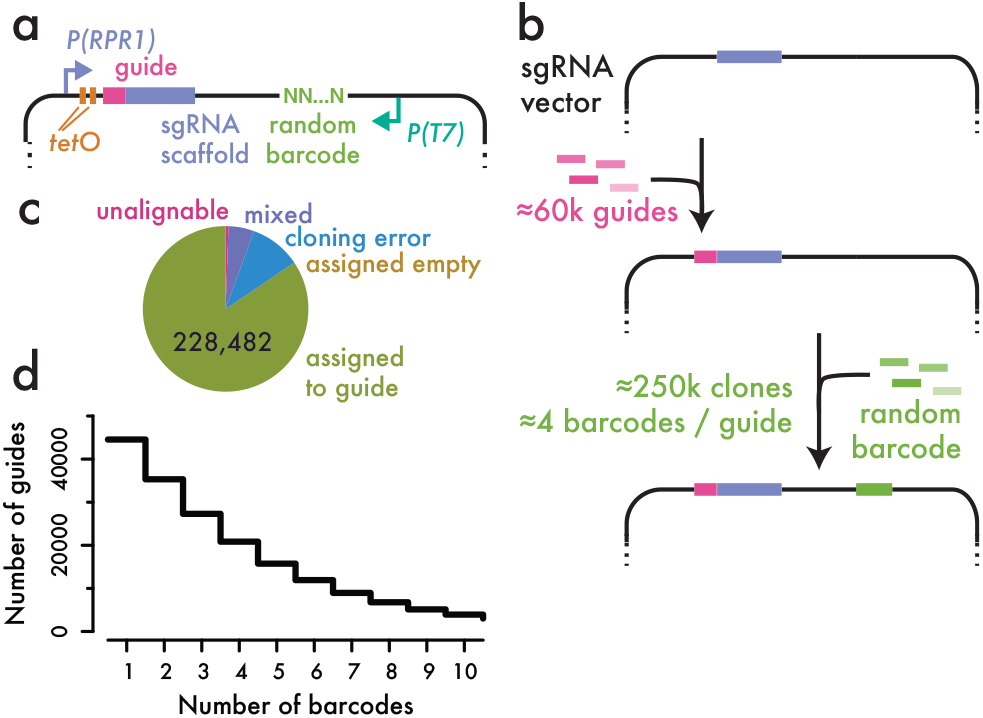
Construction of a barcoded library for inducible guide RNA expression. **(a)** Schematic of the guide RNA expression library. The *RPR1* promoter is regulated by two *tetO* operator sites. In the presence of tetracycline, this promoter is de-repressed and drives expression of a variable guide RNA sequence in a constant sgRNA scaffold. A random nucleotide barcode with an adjacent T7 RNA polymerase promoter is embedded elsewhere in the plasmid. **(b)** Schematic of the process for generating the guide RNA library. Guides are cloned first, and then barcodes are added in a transformation with controlled diversity. **(c)** Distribution of barcode-to-guide assignment results, illustrating the high frequency of errors in cloned guide RNAs. **(d)** Cumulative distribution of the number of barcodes assigned to each guide RNA.

We linked each barcode to its associated guide by high-throughput sequencing. In order to ensure reliable guide RNA assignments, we required at least three independent sequencing reads to establish a barcode-to-guide assignment. We identified ~270,000 barcodes, in good agreement with our expectation for ~250,000 distinct clones in the library. We excluded ~10% of barcodes that were linked to guides with errors introduced in cloning and synthesis (Fig. 3c). The high rate of defective guides emphasizes the value of barcoded libraries, which can identify these ineffective constructs. We also eliminated barcodes with substantial evidence linking them to two distinct guide RNAs (~5% of the total). Our library included ~45,000 distinct guides, with a median of 3 barcodes per guide and ~35,000 guides linked to more than one barcode (Fig. 3d). We also recovered 344 distinct barcodes (~1% of the total) lacking a guide RNA entirely and thus expressing only the truncated single guide RNA scaffold. We presume that these “empty” guide RNA expression constructs will have little phenotypic effect and treat these barcodes as internal negative controls.

### CRISPRi growth phenotypes recapitulate known loss-of-function phenotypes genome-wide

We wished to assess the growth phenotypes of our CRISPRi guides in a pooled yeast population. Plasmids containing guides that slow cell growth will decrease in abundance because they replicate along with the host cell, and we can measure the depletion of the associated barcodes by high-throughput sequencing. We wanted to ensure that even guides with strong negative phenotypes were present in our population at the start of the experiment, however. By using an inducible promoter to drive guide RNA expression (Farzadfard et al., 2013; Smith et al., 2016), we were able to establish a pooled population of cells that contain a diverse library of guide RNA plasmids, but do not express these guides (Fig. 4a, 4b). We then induced guide expression and followed the changes in the abundance of each guide, driven by its CRISPRi phenotype (Fig. 4a, 4c).

**Figure 4.**
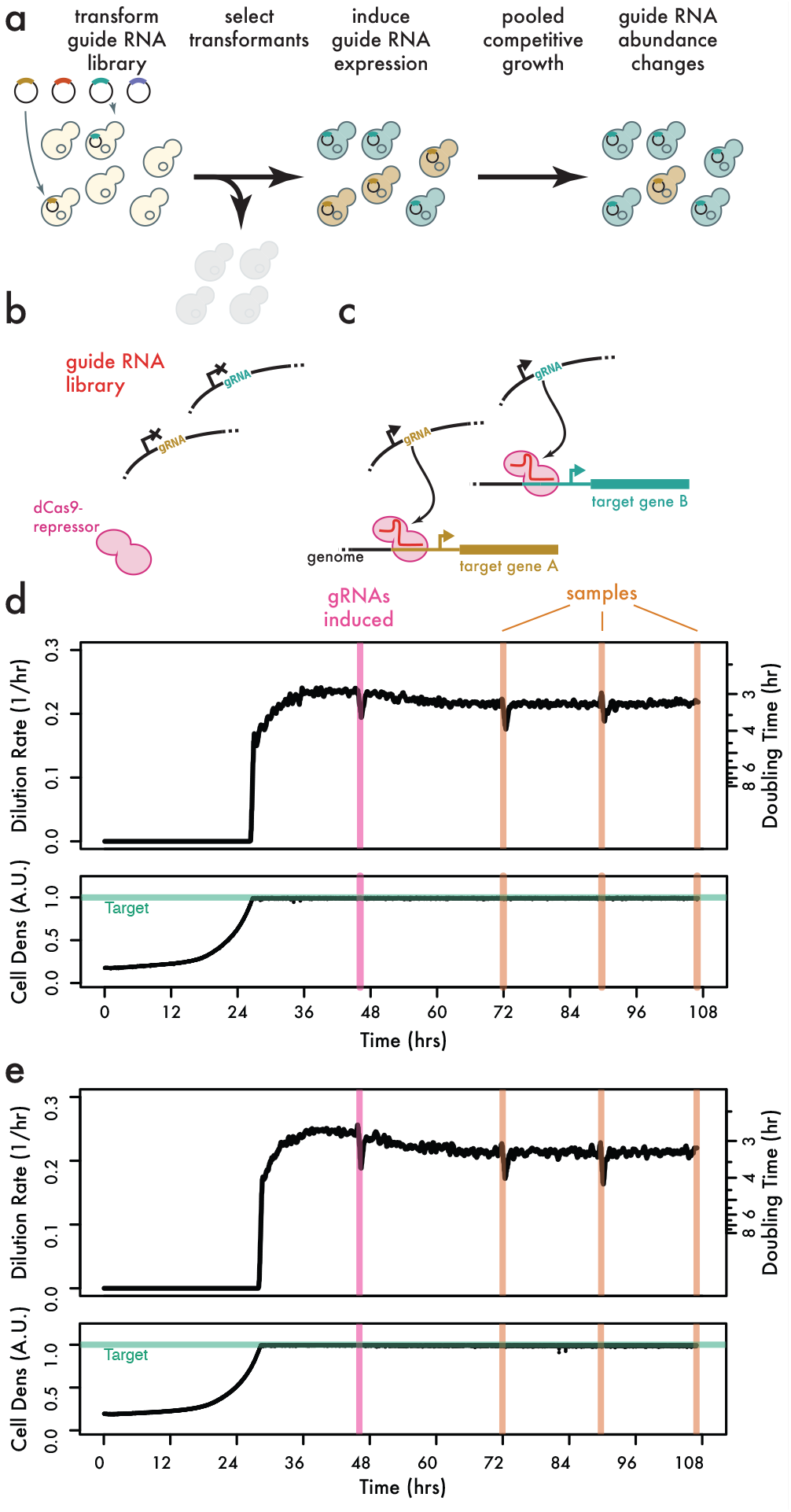
Pooled competitive growth of a diverse guide RNA library. **(a)** Schematic of the competitive growth experiment. **(b)** Cells expressing the dCas9-Mxi effector protein are transformed with guide RNA expression plasmids and selected under non-inducing conditions. **(c)** Upon guide induction, dCas9-Mxi binds target gene promoters and reduces transcription. **(d, e)** Replicate competitive growth experiments. Dilution rate corresponds to growth rate after cultures reach the target cell density. Timepoints for guide RNA induction and initial sampling is shown, along with subsequent sampling timepoints.

We also sought to maintain consistent culture conditions during the course of our competitive pooled growth experiment. After transforming our guide RNA expression library into yeast, we selected transformants — without guide induction — by growth in continuous liquid culture using a turbidostat bioreactor (McGeachy et al, 2019). We then used this selected population to inoculate a second bioreactor culture in yeast minimal media. After the cultures achieved a consistent growth rate in minimal media, we sampled the population and added tetracycline to induce guide RNA expression (Fig. 4d, 4e). We then took three additional samples over ~60 hours of growth in the presence of tetracycline and prepared high-throughput sequencing libraries to quantify barcode abundance at each timepoint and in each replicate. We further prepared replicate samples from each culture at the final timepoint in order to obtain an empirical estimate of the technical variability in our barcode abundance measurements.

Barcode abundances followed exponential dynamics during competitive growth, reflecting the fitness of the associated guide RNA. For example, the barcodes linked with one individual guide targeting *SUI3*, an essential gene encoding a translation initiation factor, declined consistently following guide induction and were almost gone after 12 generations (Fig. 5a). The rate of decline was similar for two distinct barcodes linked to this guide in each of the two replicate cultures, demonstrating that barcode abundance changes provide a robust and quantitative measure of fitness. Likewise, two distinct barcodes for a guide targeting *STV1*, a non-essential gene, show a reproducible but more gradual decline in abundance (Fig. 5b). These individual trajectories suggested that we could model these barcode sequencing data and infer quantitative growth rates.

**Figure 5.**
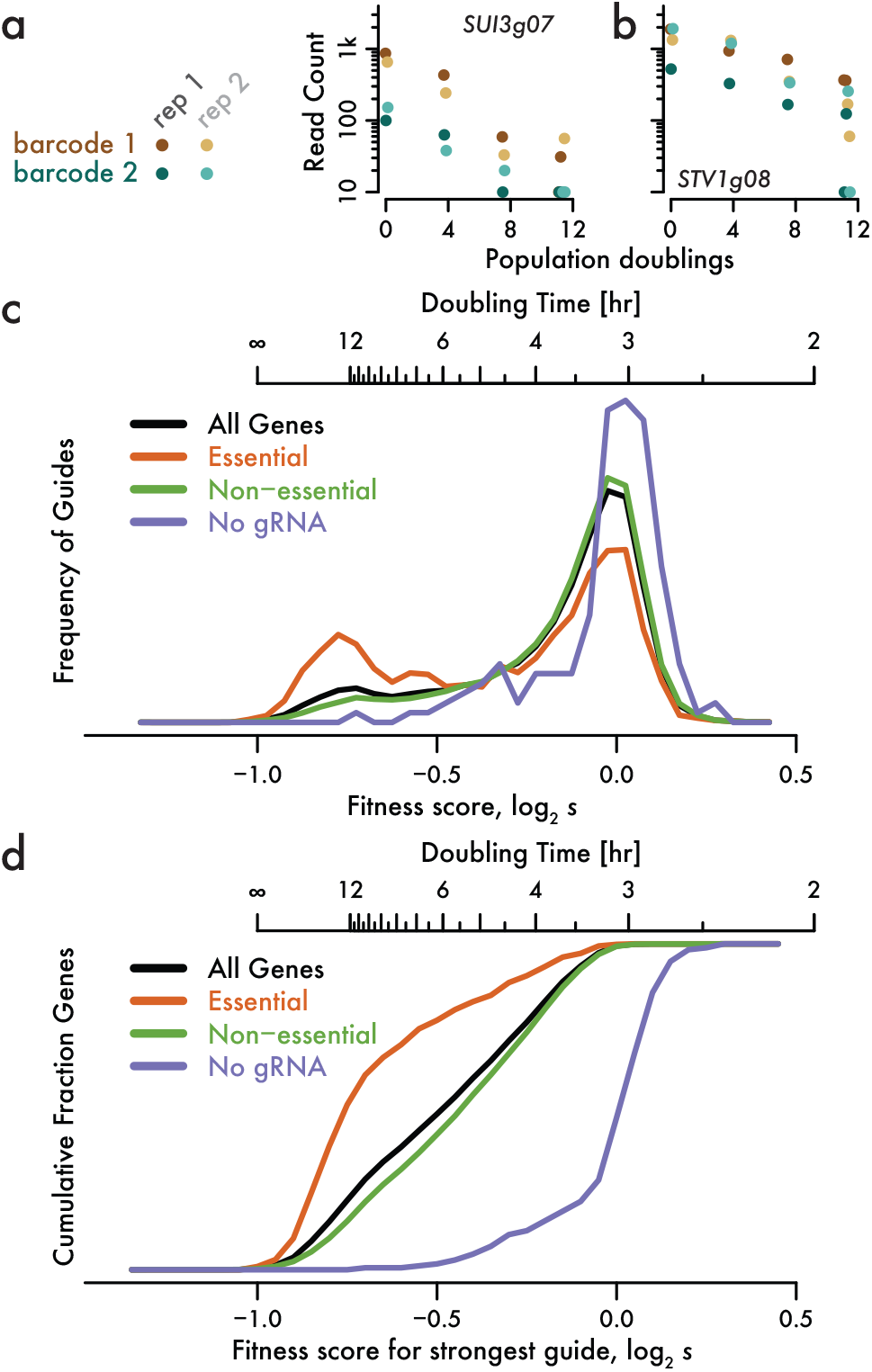
Inferring fitness effects from guide RNA abundance changes. **(a, b)** Consistent rate of exponential decay in abundance for guides targeting (a) *SUI3*, which encodes eIF2β, and (b) *STV1* during pooled, competitive growth. Two distinct barcodes are shown for each guide in two replicate cultures. **(c)** Enrichment of strong negative fitness effects in guides targeting essential genes. Guide-level fitness estimates are shown for all unambiguous promoters, as well as classification according to essentiality. Barcode-level analysis is shown for all barcodes linked to non-targeting guides. **(d)** Most essential genes have at least one strongly deleterious guide. The most negative fitness effect across all guides is shown for genes with unambiguous promoters. The cumulative distribution of barcode fitness effects is shown for non-targeting barcodes.

We took this approach to determine the fitness effect of 35,223 guide RNAs. A substantial subset of guides showed a strong negative fitness, while we saw no strong positive effects, consistent with our expectation that gene knock-down is much more likely to slow growth than to accelerate it (Fig. 5c). As we knew the number of generations between each sample, we could calibrate these measurements and obtain an actual selective coefficient s reflecting the change in abundance over one doubling of the overall population. We use the fitness score log_2_ *s*, where a fitness score of 0 corresponds to doubling at the same rate as the population overall, and a cell that ceases growth entirely has *s* = 0.5 and a fitness score of −1. We saw many fitness scores around −0.8 or below, but essentially none below −1. The minimum fitness we observe does not reflect any inherent limitation of our model, which could capture the dynamics of a guide whose abundance declined faster than 2-fold each generation. We interpret this lower bound as an indication that our fitness estimates are quantitatively accurate. Our experiment measures directly the abundance of guide expression plasmids, which are likely to persist in genetically eliminated cells that could never again divide, and even in the cell wall “ghosts” remaining when yeast lose plasma membrane integrity. Such a persistent, non-replicating plasmid will decline in relative abundance by half each generation, yielding a fitness of −1.

We expected that strongly negative fitness effects would arise most often in guide RNAs that block expression of essential genes. To avoid ambiguity in determining the gene targeted by a guide RNA, we excluded divergent promoters where CRISPRi could in theory affect either or both genes (Fig. 1d) and restricted our analysis to 3,521 unambiguous genes. Among this group, guides targeting one of 644 essential genes showed a clear bimodal fitness distribution with a distinct peak at very low fitness (Fig. 5c). In contrast, non-essential genes showed a modest depletion at very low fitness. Even in carefully designed guide RNA libraries, the majority of guides are ineffective, and so it is expected that many guides targeting essential genes nonetheless show little or no growth defect. Importantly, however, our library contained at least one guide with a strong fitness effect for almost every essential gene (Fig. 5d). This result gave us confidence that our library contained effective guides against most genes. Among non-essential genes, we saw that guides provoking a serious fitness defect (log_2_ *s* < −0.5) were strongly enriched for ribosomal proteins, as revealed by significant (*q* < 0.01) results in gene ontology analysis. Many yeast ribosomal proteins are duplicates, and so they are not individually essential but show significant growth phenotypes when deleted (Cheng et al, 2019).

### Guide RNA position predicts guide efficacy, clarifying assignments of guides to target genes

Our guide RNA library includes effective guides against most essential genes, along with many ineffective guides for these same targets. This compendium offered the opportunity to better understand what features predicted guide efficacy. Focusing on the 644 essential genes with unambiguous promoters, we analyzed 1,967 guides that targeted one of these genes and had at least two distinct, high-abundance barcodes in our data set.

The position of the guide relative to the transcription start site greatly impacted the efficacy of CRISPRi (Fig. 6a). All of our guides fell within a 240 nucleotide window around the transcription start site, and we ensured that guides were distributed across this region at each promoter. Strong fitness effects were most likely for guides binding roughly 50 bases upstream of the transcription start site, and fell off substantially on either side. Furthermore, we saw differences between guides matching the coding versus the template strand (Fig. 6b). On either strand, guides showed the strongest average fitness effect when the invariant protospacer-adjacent motif (PAM) recognized directly by the Cas9 protein fell 50 bases upstream of the transcription start site.

**Figure 6.**
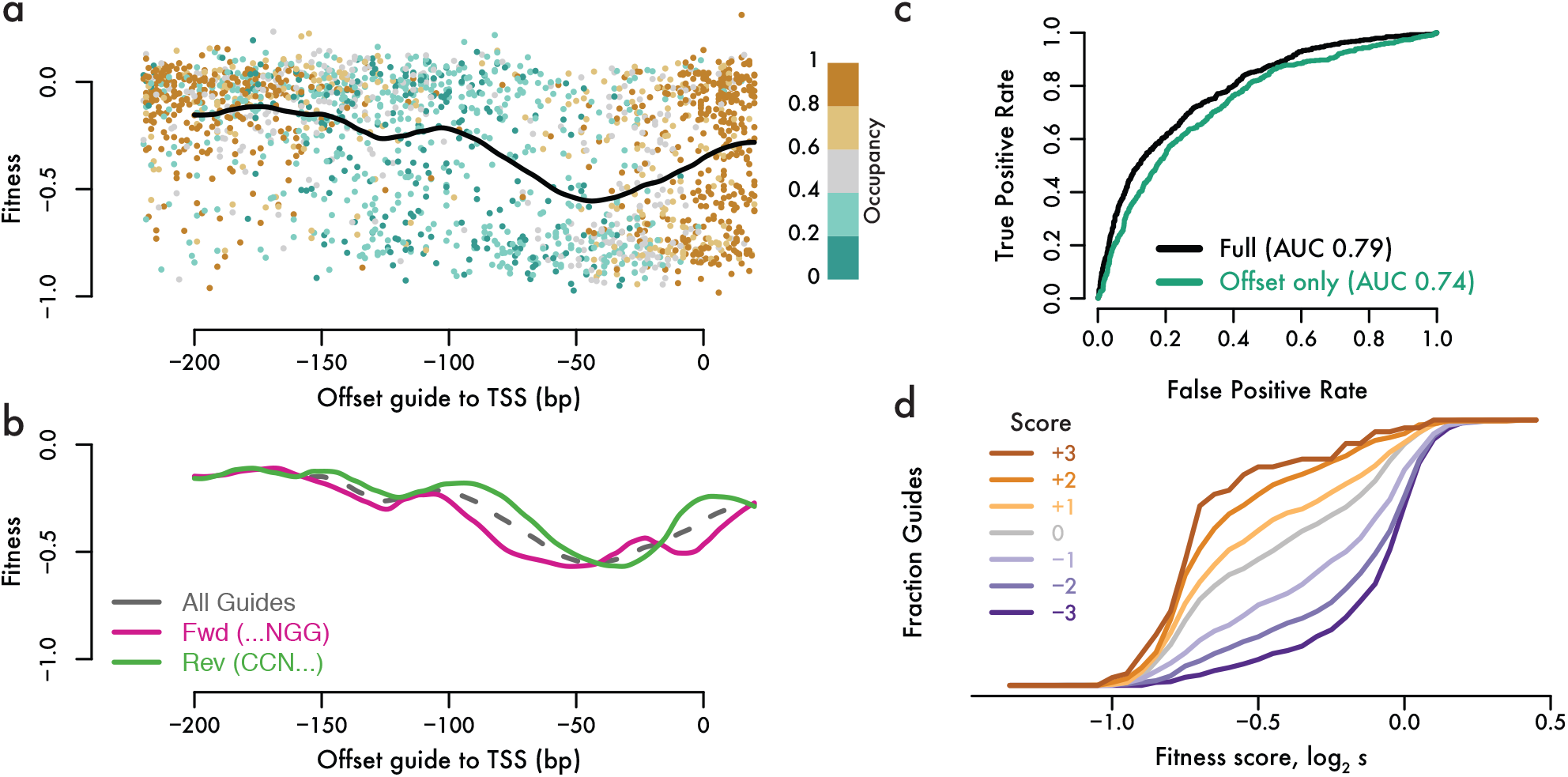
Accurate predictions of guide activity. **(a)** Highly active guides against essential genes bind ~50 bp upstream of the transcription start site. The fitness effect produced by guides targeting essential genes serves as a proxy for their activity. Individual guides targeting non-divergent essential genes with at least two independent barcodes are plotted according to their position, fitness effect, and absolute chromatin occupancy as determined by DNA methylation accessibility, along with a local regression of fitness effect against position. **(b)** Guide activity varies according to the guide strand, as seen in local regressions for guides on the same (Fwd) or opposite (Rev) strand as the target gene. **(c)** Receiver operating characteristic for logistic regression models of guide activity. The full model includes methylation-based accessibility data (ODM-Seq) and guide sequence along with the position of the guide relative to the transcription start site. **(d)** Guide score predicts fitness effects in a held-out test set of guides targeting essential genes.

We used the strong positional bias of CRISPRi efficiency to resolve the likely target genes for guides at divergent promoters. Transcriptional start sites are typically separated by over 200 nucleotides, and so very few guides fall into the high-efficacy region for both genes. At a few closely-spaced promoters, it may prove impossible to inhibit just one gene potently and specifically by CRISPRi. In most cases, however, we can determine the target gene for each guide, increasing the number of specific guides for each gene in our library (Fig. 1c).

Accessibility of target site DNA correlates with guide activity in yeast and mammalian CRISPRi. Chromatin accessibility is partly confounded with position effects, as active yeast promoters show well-defined organization, with a nucleosome-free region around the transcription start site bounded by a positioned +1 nucleosome (Yuan et al, 2005). To investigate this effect in our library, we took accessibility measurements from a recent study that probed for DNA sensitive to in vitro methylation (ODM-Seq), which is blocked by nucleosome occupancy (Oberbeckmann et al, 2019). While the correlation between accessibility and position is apparent, it does not seem to explain the pattern of guide activity. Open chromatin typically extends over 100 bp upstream of the transcription start site, whereas guide activity falls off at shorter distances (Fig. 6a). Thus, it appears that position and accessibility contribute separately to guide activity, leading us to seek a statistical model that could predict effective CRISPRi.

We developed a logistic regression model for active guides based on the target site position, sequence, and accessibility. We accounted for the complex relationship between position and activity empirically, using the local regression of quantitative fitness effect against the distance to the transcription start site (Fig. 6a) as one parameter in a larger regression model. Position alone predicted activity well (AUC 0.74; Fig. 6c), and performance improved substantially when methylation-based ODM-Seq accessibility and nucleotide sequence features were added (AUC 0.79; Fig. 6c). Both accessibility and sequence features individually improved model performance in *k*-fold cross-validation (Figure S3). Incorporating strand-specific position effects (Fig. 6b) decreased residual variance significantly, but did not improve model performance in cross-validation (Figure S3), and so we retained the strand-independent model. We also tested the relative contributions of ODM-Seq and ATAC-Seq accessibility data. While ATAC-Seq alone did contribute to model performance, we found that ODM-Seq yielded significant further improvement, whereas adding ATAC-Seq data did not improve on a model that already incorporated ODM-Seq data, and so we used ODM-Seq accessibility alone.

Our model produced well-calibrated predictions of guide activity when assessed on a separate set of 3,480 guides. These guides were held out of model development and validation, either because they targeted essential genes at divergent promoters, or because they were linked with only one high-abundance barcode. Most high-scoring guides produced fitness effects, whereas few low-scoring guides impaired growth, demonstrating that our regression model generalized well to these other guides (Fig. 6d). Furthermore, among guides with a logit-transformed score of 0, corresponding to equal odds of activity or inactivity, we observed a median fitness close to our threshold value for active guides. This quantitative agreement between model predictions and measured guide activity indicates that our score directly indicates the likelihood that a guide is active.

## DISCUSSION

We provide a strategy for genome-wide CRISPR interference screening in budding yeast. We overcome the unpredictable activity of guide RNAs by designing ten distinct guides per gene, producing a library that contains at least one active guide against most genes, as assessed by our ability to induce growth defects on essential targets (Fig. 5d). We also show that random nucleotide barcodes with linear IVT-RT amplification provide significant advantages for robust and quantitative CRISPR screening. Random nucleotide barcodes enable multiple, independent measurements of guide RNAs within a single experiment (Fig. 5a, 5b) and distinguish sequencing errors from defective guides (Fig. 3c). Linear amplification by in vitro transcription improves the quantitative reproducibility of barcode abundance measurements in sequencing data and reduces the occurrence of extreme outliers (Fig. 2). Barcoding and linear amplification are complementary and separable features — barcodes can be amplified by PCR, of course, and an embedded T7 RNA polymerase promoter can be used to transcribe guides themselves in vitro.

Systematic genetics in budding yeast benefits from a wealth of techniques for analyzing defined loss-of-function phenotypes. Most notably, barcoded deletion strains are available for most non-essential genes (Giaever et al., 2002), and resources such as partial loss-of-function “DAmP” alleles (Schuldiner et al, 2005) and titratable promoter alleles (Mnaimneh et al, 2004) are available for many essential genes. Inducible CRISPRi offers distinctive advantages relative to these and other existing resources. It treats essential and non-essential genes on an equal footing by providing consistent partial-loss-of-function alleles resulting from reduced transcription. It also avoids the accumulation of suppressor mutations and other genetic aberrations that arise during long-term propagation of strains with heritable genetic lesions.

Genome-wide CRISPRi also provides practical features that are advantageous for many comprehensive genetic analyses. It is straightforward to carry out screens in nearly any genetic background, as guides are introduced in a single, pooled transformation of episomal plasmids. In contrast, other approaches for genome-wide genetic analysis depend on elegant but complex mating and selection schemes to recombine genetic lesions into the desired haploid genotype (Pan et al, 2004; Tong et al, 2001). Indeed, the ease of carrying out a screen using this CRISPRi library approaches that of standard forward genetic screens. Deep sequencing of guide-linked barcodes (or guides themselves) then reveals a quantitative profile of phenotypes across the genome. We anticipate many uses for this screening system, as well as future refinements of guide RNA design based on data presented here.

## MATERIALS AND METHODS

### Plasmids

Plasmids were constructed by standard molecular biology techniques as described below and verified by Sanger sequencing (Table S1). Restriction enzymes were obtained from NEB and high-fidelity (HF) variants were used when available. Q5 polymerase (NEB M0491S) was used for PCR, assembly reactions were carried out using Gibson Assembly Master Mix (NEB E2611L).

<u>pNTI647</u> was generated by amplifying the adjacent dCas9-Mxi and TetR expression cassettes from pNTI601 (pRS416-dCas9-Mxi1 + TetR + pRPR1(TetO)-NotI-gRNA, Addgene #73796) (Smith et al., 2016) using primers NM721 and NM734 (Table S1). This insert was assembled into pCfB2225 (AddGene #67553), an “EasyClone 2.0” vector for KanMX-marked integration into the XII-2 safe harbor location (Stovicek et al, 2015).

<u>pNTI661</u> was generated in several steps from pNTI601. The *URA3* marker was replaced by the *K. lactis LEU2* marker from pUG73 (Gueldener et al, 2002) by amplifying this marker using primers NI-993 and NI-994 (Table S1), as well as amplifying a backbone fragment of pNTI601 using primers NI-995 and NI-996 (Table S1), and assembling these back into pNTI601 digested with SpeI and KpnI. Primers KS524 and KS525 (Table S1) were used to amplify the region of the vector excluding dCas9-Mxi1 and TetR, which was recircularized by Gibson assembly. The barcode site was introduced by amplifying the guide RNA expression cassette with NI-1019 and NI-1020 and re-ligating the resulting product back into the vector after a SacI/SpeI digestion of both vector and PCR amplicon. Finally, the NotI site for guide RNA cloning was replaced with a BamHI-HindIII cassette by digesting the vector with NotI and performing Gibson assembly with the NI-1030 oligonucleotide.

<u>pNTI698</u> was generated by amplifying the *HIS3*, *MET17*, and *URA3* genes from pHLUMv2 (AddGene #64166) (Mulleder et al, 2016) using p698Fwd and p698Rev primers (Table S1). This insert was assembled into pCfB2223 (AddGene #67544), an “EasyClone 2.0” vector for KanMX-marked integration into the X-3 safe harbor location (Stovicek et al., 2015), digested with EcoNI. Note that the KanMX marker is disrupted by the *HIS3-MET17-URA3* cassette and the plasmid no longer confers resistance.

### Yeast

#### Strains

Yeast were derived from *S. cerevisiae* strain BY4741 (ThermoFisher), a haploid MATa his3Δ1 leu2Δ0 LYS2 met15Δ ura3Δ0 derivative of S288c.

<u>NIY416</u> was derived from BY4741 by transformation with integrating plasmid pNTI647 digested with NotI, followed by selection for kanamycin resistance.

<u>NIY425</u> was derived from NIY416 by transformation with integrating plasmid pNTI698 digested with NotI, followed by selection for Ura and Met prototrophy.

#### Media

Minimal media was prepared using 67. g / l yeast nitrogen base with ammonium sulfate and without amino acids (BD 291920) and 200. g / l dextrose (Fisher D16-500). Synthetic complete drop-out media minus leucine (SCD -Leu) was prepared using 67. g / l yeast nitrogen base with ammonium sulfate and without amino acids, 1.62 g / l synthetic drop-out mix minus leucine (US Bio D9626), and 200. g / l dextrose.

#### High-efficiency transformations

High-efficiency yeast transformations were carried out by growing yeast cultures overnight at 30 °C with shaking and diluting these cultures to prepare fresh dilution cultures at an OD600 of 0.05. Dilution cultures were grown at 30 °C with shaking until they reached an OD600 of 0.5 and then 20 ml of culture was taken for each transformation. Cells were pelleted by centrifugation at 3,000 × g for 10 minutes and the supernatant was decanted. Cells were resuspended in 10. ml sterile deionized water and pelleted again by centrifugation at 3,000 × g for 5 minutes, the supernatant was decanted, and any residual liquid was removed with a pipettor. Cells were then resuspended in 1.0 ml lithium acetate 100 mM, transferred to a microcentrifuge tube, and pelleted by centrifugation at 10,000 × g for 10 seconds. Supernatant was removed by aspiration and cells were resuspended in 1.0 ml lithium acetate 100 mM and pelleted again at 10,000 × g for 10 seconds. Supernatant was removed by aspiration, and 240 *μ*l of 50% w/v polyethylene glycol was layered gently on top of cells, followed by 20. *μ*l of freshly boiled salmon sperm DNA 10 mg / ml (Invitrogen 15632011), 36. *μ*l lithium acetate 1.0 M, and 64. *μ*l plasmid DNA. The microcentrifuge tubes were then vortexed vigorously to resuspend cells and incubated for 20 minutes in a 42 °C water bath, vortexing once during the incubation to maintain cells in suspension. Following this incubation, cells were pelleted by centrifugation at 10,000 × g for 10 seconds and the transformation mixture was removed with a pipettor. Cells were resuspended in 1 ml sterile deionized water, pelleted by centrifugation at 10,000 × g for 10 seconds, and the water was removed with a pipettor. Finally, cells were resuspended in 1.0 ml sterile deionized water per transformation.

### Guide library design

#### External data sets

Yeast genome sequence (R64-1-1, sacCer3) (Engel et al, 2014) and CDS annotations (Cherry et al., 2012) were downloaded from the UCSC genome browser, and yeast gene information was downloaded directly from the Saccharomyces Genome Database (Cherry et al., 2012). Transcript isoform data was obtained from Pelechano et al. (Pelechano et al., 2013). ATAC-seq data was obtained from from Schep et al (Schep et al., 2015), GEO accession GSE66386.

#### Gene annotations

All major transcript isoforms (mTIFs) from Pelechano et al. (Pelechano et al., 2013)annotated to cover one intact ORF were considered for gene annotation. Considering the set of mTIFs for a gene, the modal (highest read count) transcription start site (TSS) was chosen as the representative transcription start site for the gene. When no transcript was annotated for the ORF, the annotated CDS was used for guide design and target prediction.

#### Guide scoring

All possible guides were identified by searching for GG dinucleotides, representing the Cas9 protospacer adjacent motif (PAM) in the yeast genome sequence. Guide site uniqueness was assessed by aligning each target sequence (20 base protospacer followed by “NGG” PAM) against the yeast genome reference using Bowtie2 [22388286]. Target sequences with multiple perfect genomic alignments were considered non-unique. Guides were associated with gene TSSes when the center of the target sequence fell between −220 and +20 nucleotides relative to the TSS. Guides were associated with CDS genes when the center of the sequence fell between −350 and 0 nucleotides relative to the CDS. Guides were considered specific when these targeting rules associated the guide with only one single target gene. Target accessibility was determined by averaging ATAC-Seq accessibility, ranging from 0.0 for inaccessible to 1.0 for fully accessible, across all nucleotide positions in the target sequence in two replicate ATAC-Seq data sets. When no data was available, a value of 0.0 was used.

#### Guide selection

Guides were prioritized by first preferring unique guides, and then specific guides, and finally by greater ATAC-Seq accessibility. For each TSS-annotated gene, the highest-scoring guides were chosen for three zones spanning [−220, −141], [−140, −61], and [−60, +20] nucleotides relative to the TSS. For each CDS-annotated gene, the highest-scoring guides were chosen for four zones spanning [−350, −271], [−270, −191], [−190, −111], and [−110, −30] nucleotides relative to the start of the CDS. Additional guides were chosen, highest score first, until ten guides were chosen or all possible guides in the targeting region were exhausted.

### Barcoded guide expression library

#### Guide library construction

The guide RNA expression vector pNTI661 was digested by taking 3.0 *μ*g plasmid in a 75. *μ*l reaction with 1x final concentration CutSmart buffer (NEB B7204S) with 60 U BamHI-HF (NEB R3136L) and 60 U HindIII-HF (NEB R3104S), incubated for 1 hour at 37 °C, and then purified with a DNA Clean & Concentrator (Zymo D4013). The guide RNA oligonucleotide library was amplified using Q5 polymerase (NEB M0491S) according to the manufacturers instructions, using 100 pg guide oligonucleotide pool (CustomArray, Inc.) as a template and oligonucleotides NM636 and NM637 (Table S1) for amplification, with 15 cycles of amplification using 10 s denaturation, 15 s annealing at 58 °C, and 15 s extension. Amplified guide RNAs were cloned in a 100 *μ*l assembly reaction with 1.0 *μ*g linearized pNTI661 and 1.7 *μ*l guide RNA PCR using 2 × NEBuilder HiFi DNA Assembly Master Mix (NEB E2621L), which was incubated for 1 hour at 50 °C and then purified with a DNA Clean & Concentrator with final elution into 10. *μ*l. Purified DNA was used to transform high efficiency competent 10-beta *E. coli* (NEB C3019H), using 2.5 *μ*l purified DNA per reaction in four independent transformations of 50 *μ*l competent cells. Following transformation, transformations were pooled into 100 ml LB Carb liquid media and grown with vigorous shaking until reaching an OD600 of 3. Plasmid DNA was extracted with a QIAGEN Plasmid Midi Kit (QIAGEN 12143).

#### Barcode addition

The guide expression library was digested again with BamHI-HF along with exonucleases in order to digest and degrade the majority of the guide-free plasmids. A 50 *μ*l digestion reaction was prepared using 2 *μ*g plasmid DNA in 1x final concentration CutSmart buffer with 20 U BamHI-HF, 5 U lambda exonuclease (NEB M0262S), and 20 U *E. coli* exonuclease I (NEB 0293S). Digestion was carried out for 1 hour at 37 °C, followed by heat inactivation for 20 minutes at 80 °C. DNA was then purified using a Zymo DNA Clean & Concentrator column, with elution into 20. *μ*l. The library was then linearized for barcode assembly in a 50 *μ*l digestion reaction using 18. *μ*l of eluted DNA from the previous digestion in 1x final concentration CutSmart buffer with 20 U SphI-HF (NEB R3182S). Digestion was carried out for 1 hour at 37 °C, and DNA was purified again using a Zymo DNA Clean & Concentrator column.

Random nucleotide barcodes with embedded T7 RNA polymerase promoters were generated by PCR amplification from 1.0 *μ*l NI-1026 oligonucleotide using NI-1027 and NI-1041 oligonucleotide primers (Table S1). A 50 *μ*l PCR using Q5 polymerase (NEB M0491S) according to the manufacturers instructions, with 15 cycles of amplification using 5 s denaturation, 10 s annealing at 65 °C, and 5 s extension. Product was purified using a DNA Clean & Concentrator column. Amplified barcodes were introduced in a 100 *μ*l NEBuilder HiFi Assembly reaction containing 1 *μ*g linearized guide library and 110 ng purified barcode PCR. DNA was purified using a DNA Clean & Concentrator column with final elution into 10 *μ*l. Purified DNA was used to transform high efficiency competent 10-beta *E. coli*, using 2.5 *μ*l purified DNA per reaction in four independent transformations of 50 *μ*l competent cells. Following transformation, transformations were pooled into a single, 4.0 ml pool. Dilutions were plated on LB Carb agar plates to assess transformation efficiency, and 55% of the transformation was used to inoculate a 50 ml LB Carb culture while 22%, 8%, 6%, and 4% were used to inoculate four separate 25 ml LB Carb cultures. Higher-inoculum 55% and 22% cultures were grown at 26 °C overnight, while lower-inoculum 8%, 6%, and 4% cultures were grown at 30 °C overnight. Based on the estimated yield of ~1.1 M transformants, the 22% culture was selected. DNA was isolated using a QIAGEN Plasmid Mini kit to produce the barcoded guide expression library.

### Comparative barcode amplification

#### Guide library transformation and yeast growth

BY4741 was transformed with barcoded guide expression library in one high-efficiency transformation of ~100 M cells using 64 *μ*l of plasmid DNA at 100 ng / *μ*l. Dilutions were plated on SCD -Leu agar plates in order to estimate the transformation efficiency, indicating a yield of ~330,000 independent transformants. The rest of the transformation was used to inoculate 100 ml of SCD -Leu media and grown for ~24 hours at 30 °C with shaking, at which point the OD600 increased roughly 4-fold, to 0.82. A new 100 ml SCD -Leu culture was inoculated with 400 *μ*l of this culture and growth at 30 °C with shaking was continued overnight to yield a final OD600 of 1.7. Four aliquots of 25 ml each were taken for yeast plasmid DNA extractions. Yeast were pelleted by centrifugation for 10 minutes at 3,100 × g, and media was discarded. Cells were resuspended in 1.0 ml sterile deionized water, pelleted 10,000 × g for 30 seconds, and water was removed by aspiration. Washed yeast pellets were stored at −80 °C.

#### Linear amplification by in vitro transcription

Half of one plasmid extraction was used to prepare a 25 *μ*l digestion in 1x final concentration CutSmart buffer with 20 U XhoI (NEB R0146L) and incubated 1 hour at 37 °C. DNA was purified using a DNA Clean & Concentrator column with elution into 20. *μ*l, and 18 *μ*l of purified DNA was used as template in a 30 *μ*l HiScribe T7 Quick High Yield RNA Synthesis reaction (NEB E2050S) following the protocol for short templates and incubated overnight at 37 °C. Template was degraded by adding 20 *μ*l water followed by 4 U DNase I and continuing incubation for 15 minutes at 37 °C and RNA was then purified using an RNA Clean & Concentrator, with final elution into 15. *μ*l. Purified RNA was assessed using a High Sensitivity RNA ScreenTape with an Agilent TapeStation 2200. Reverse transcription was carried out using 10 ng of purified RNA in a reaction with ProtoScript II (NEB M0368S) using 2.0 pmol NI-1032 as a gene-specific primer (Table S1). Primer and template were denatured 5 min at 65 °C, kept on ice to prepare reactions, and then incubated 1 hour at 42 °C followed by heat inactivation at 65 °C for 20 minutes. A 50 *μ*l PCR reaction using Q5 was prepared using 5.0 *μ*l RT product as a template without further purification, along with NEBNext Multiplex Oligos for Illumina (NEB E7600S) as primers, and amplified for 7 cycles using 5 s denaturation, 10 s annealing at 65 °C, and 10 s extension. PCR products were purified using AMpure XP beads according to the manufacturer’s instructions, using a 2 beads : 1 PCR ratio and final elution in 20. *μ*l Tris•Cl 10 mM, pH 8.0. Products were validated using a High Sensitivity D1000 ScreenTape on an Agilent TapeStation 2200, pooled, and analyzed by 50 base single-read deep sequencing on an Illumina HiSeq with 10% phiX control. Note that the first 25 bases comprise high-diversity barcode libraries whereas the subsequent bases are monotemplate.

#### Exponential PCR amplification

First-round PCR was performed using Q5 polymerase, 10% of extracted yeast plasmid DNA as a template, and primers NI-956 and NI-1032 (Table S1), and amplified for 16 cycles using 10 s denaturation, 15 s annealing at 65 °C, and 10 s extension. PCR products were purified using AMpure XP beads according to the manufacturer’s instructions, using a 2 beads : 1 PCR ratio and final elution in 20. *μ*l Tris•Cl 10 mM, pH 8.0. Second-round PCR was performed exactly as described for linear amplification by in vitro transcription, except that 1.0 *μ*l of purified first-round PCR product was used as a template. PCR libraries were validated, pooled, and sequenced in parallel with linear amplification libraries.

#### Barcode analysis

Barcode sequencing data was analyzed by trimming the 3’ adapter sequence “GCATGCGTGAAGTGGCGCGCCTGATA” using Cutadapt, discarding all sequences that either lacked a linker or contained a barcode sequence less than 10 nucleotides long. Barcodes were tabulated using a custom tool, “bc-count”, that collapses single-nucleotide mismatches. Barcode counts were collated across all four libraries and filtered to remove barcodes that occurred in only one library or had fewer than 33 reads total across all 4 libraries. Barcodes were also filtered to remove sequences containing XhoI sites. Barcode counts were plotted, and DESeq2 was used to estimate read count-dispersion relationships from barcode count tables.

### Barcoded-to-guide assignment

#### Sequencing library construction

First-round PCR was carried out in 50 *μ*l using Q5 polymerase with 100 ng barcoded guide library as template and primers NI-1038 and NI-956 (Table S1), and 12 cycles of amplification were performed using 10 s denaturation, 15 s annealing at 67 °C, and 20 s extension. PCR products were purified using AMpure XP beads at an 0.8 beads : 1 PCR ratio and final elution in 15. *μ*l Tris•Cl 10 mM, pH 8.0. Second-round PCR was performed with 1.0 *μ*l of first-round PCR as template and primers NI-798 and NI-826 (Table S1), and 15 cycles of amplification were performed using 10 s denaturation, 15 s annealing at 65 °C, and 20 s extension. PCR products were again purified using AMpure XP beads and validated using a High Sensitivity D1000 ScreenTape on an Agilent TapeStation 2200 prior to 150 base paired-end sequencing on an Illumina MiSeq. PhiX control DNA was mixed to account for monotemplate regions of the library. Barcode sequencing data is available under accession SRR10356224.

#### Sequencing data analysis

Barcodes in R1 reads were trimmed to remove the 3’ adapter sequence “GCATGCGTGAAGTGGCGCGCCTGATAGCTCGTTTAAACTG” and read pairs lacking this adapter in the R1 read, or reads with residual barcodes less than 12 nucleotides long, were discarded. Trimmed barcodes were collapsed to combine barcodes with single-nucleotide mismatches using the custom “bc-seqs” program, and guide sequences in R2 were then trimmed to remove the 5’ adapter “CGAAAC” and the 3’ adapter “AAGTTAAAAT”, leaving 20 bases of constant sequence on each side of the variable 20 nucleotide guide sequence. Read pairs where less than 20 nucleotides of residual guide sequence remained were discarded. Remaining guide sequences were aligned against a library of guide sequences using bowtie2. These alignments were used to compute barcode assignments using the custom “bc-grna” program. This tool grouped all guide alignments associated with the same barcode sequence, discarded sequences with low-quality (Q < 30) bases, and then eliminated all barcodes that lacked at least 3 high-quality guide reads. Barcodes are assigned to guides when they are supported by at least 3 high-quality reads, at least 90% of these reads align to the same guide sequence and the majority alignment to that guide has no mismatches, insertions, or deletions. Barcodes where <90% of all reads aligned to a single majority guide were considered heterogeneous and discarded. Barcodes where the majority alignment contained mismatches, insertions, or deletions were considered defective guides. The number of barcodes in each of these categories is tabulated in the “grna-assign-barcode-fates.txt” file and the high-quality barcode-to-guide assignments are given in the “grna-assign-barcode-grna-good.txt” file.

### Competitive growth

#### Guide library transformation

Guide RNA library was transformed into NIY425 as described in “High-efficiency transformations.” Three independent transformations were pooled and used to inoculate a turbidostat (McGeachy et al., 2019) containing ~200 ml SCD -Leu media at an initial OD600 of 0.1. The culture was maintained for ~24 hours at a target OD600 of 0.5, at 30 °C with continuous aeration and stirring. A 40 ml culture was combined with 40 ml fresh, pre-warmed SCD -Leu media and grown in batch culture at 30 °C with shaking for 4.5 hours, reaching an OD600 of 2.0. Cells were pelleted by centrifugation for 10 minutes at 3,100 × g, room temperature and media was discarded. Cells were resuspended in 8.0 ml sterile deionized water and split into 8 aliquots of 1.0 ml. Cells were pelleted 10,000 × g for 30 seconds and water was removed by aspiration. Cells were resuspended in 0.80 ml sterile 30% glycerol in deionized water, flash frozen in liquid nitrogen, and stored at −80 °C.

#### Competitive growth

Two independent turbidostats (McGeachy et al., 2019) each containing ~200 ml minimal media were inoculated with aliquots of the guide library transformant pool, yielding an initial OD600 of 0.1. Turbidostats were grown at 30 °C with continuous aeration and stirring, with a target OD600 of 0.5. After ~46 hours, a 50 ml sample was withdrawn from each turbidostat and processed as described for “Guide library transformation and yeast growth” in “Comparative barcode amplification. Turbidostat media was then replaced with minimal media containing 250 *μ*g / l anhydrotetracycline and growth was continued, with additional 50 ml samples taken at ~72 hours, ~90 hours, and ~107 hours.

#### Barcode abundance library construction

Plasmid DNA was extracted from frozen yeast pellets. Barcodes were amplified and sequenced as described above for “Linear amplification by in vitro transcription” in “Comparative barcode amplification,” except that 1 - 10 ng of in vitro transcription product was used as a reverse transcription template, and 12 cycles of PCR amplification were carried out in the final step of library generation.

#### Sequencing data analysis

Barcode abundance was tabulated as described above for “Comparative barcode amplification” and barcodes were matched to guides using the results of “Barcode-to-guide assignment.”

#### Fitness effect analyses

Barcodes were filtered to eliminate entries that did not have at least 64 reads tabulated for the pre-induction sample in at least one replicate culture. These filtered barcode counts were then analyzed using DESeq2 with the model *counts* ~ *gens* + *culture*, where *gens* was a numerical factor that was 0.0 for pre-induction samples and then 3.75, 7.5, and 11.25 for the three post-induction timepoints, and *culture* was a discrete factor for the two replicate cultures. The *gens* parameter from this linear model was taken as an estimate of the selective coefficient per population doubling for each barcode. Guide-level analysis was performed by taking the weighted mean of the estimate for each individual barcode, using the standard error estimate to compute 1/Var weights for each barcode. Fitness effect distributions were calculated by first filtering for genes with unambiguous guide targeting, where TSS data was available and no guide RNA had an alternate target gene identified by our approach. A list of essential genes was downloaded from the Saccharomyces genome deletion project (Giaever et al., 2002; Winzeler et al., 1999).

#### Guide efficacy analysis

Efficacy models were fitted using 1,967 guides against essential genes with unambiguous targeting and fitness effects derived from more than one barcode. The offset between the guide target and the transcription start site was calculated based on the center of the 23 nucleotide target sequence. The relationship between fitness effect and guide-to-TSS offset was modeled with a local regression (α = 0.25) across the −220 to +20 range used for guide selection. Accessibility data was derived from Oberbeckmann et al. ODM-Seq data (Oberbeckmann et al., 2019), using the lowest occupancy value in a 33 nucleotide window including the full target sequence and 5 flanking nucleotides on each side. The fitness threshold for active guides, *s* < −0.38 was defined according to the 5th percentile of all negative controls. Logistic regression against activity classification was performed using the model *active ~ OffsetPred + ODM + nt01 + … + nt20*, where *OffsetPred* was the predicted value from the local regression of guide position, *ODM* was the ODM-Seq accessibility data, and *nt01* through *nt20* were twenty discrete factors representing the variable guide sequence. Alternative models excluded the *ODM* variable or the twenty sequence factors, included ATAC-Seq data from Schep et al (Schep et al., 2015) used in guide design, or used two distinct, strand-specific local regressions for *OffsetPred*. Models (local regression and logistic regression together) were tested by *k*-fold cross-validation with *k* = 10, and the final model was generated using all guides. This final model was used to score 3,480 guides (1,491 active, i.e., log_2_ *s* < −0.38) against essential genes that had been held out of the model development because they targeted divergent promoters or had just one barcode quantified.

## ACKNOWLEDGEMENTS

This work was supported by grants DP2CA195768 and R01GM130996 from the National Institutes of Health (N.T.I.). pRS416-dCas9-Mxi1 + TetR + pRPR1(TetO)-NotI-gRNA was a gift from Ronald Davis. pCfB2223 and pCfB2225 were gifts from Irina Borodina. pHLUM (version 2) was a gift from Markus Ralser.

## AUTHOR CONTRIBUTIONS

NJM and NTI conceived and designed the study. NJM, ZAM, and KS constructed and validated the CRISPRi plasmids. NJM, ZAM, and RM tested barcode amplification strategies. ZAM carried out the growth screen with assistance from RB. NTI analyzed sequencing data. All authors read and approved the final manuscript.

## DATA AVAILABILITY

The datasets and computer code produced in this study are available in the following databases:

- High-throughput sequencing data are available from the NCBI SRA under BioProject PRJNA579997.
- Scripts to analyze these data and generate figures for this manuscript, along with key data tables including the guide RNA library, the barcode-to-guide assignment table, and the guide-level fitness data, are available on GitHub at https://github.com/ingolia-lab/yeast-crispri.
- Plasmids and barcoded guide RNA libraries are available from AddGene.
- General information is available at https://ingolia-lab.github.io/yeast-crispri/

---

**Figure S1.**
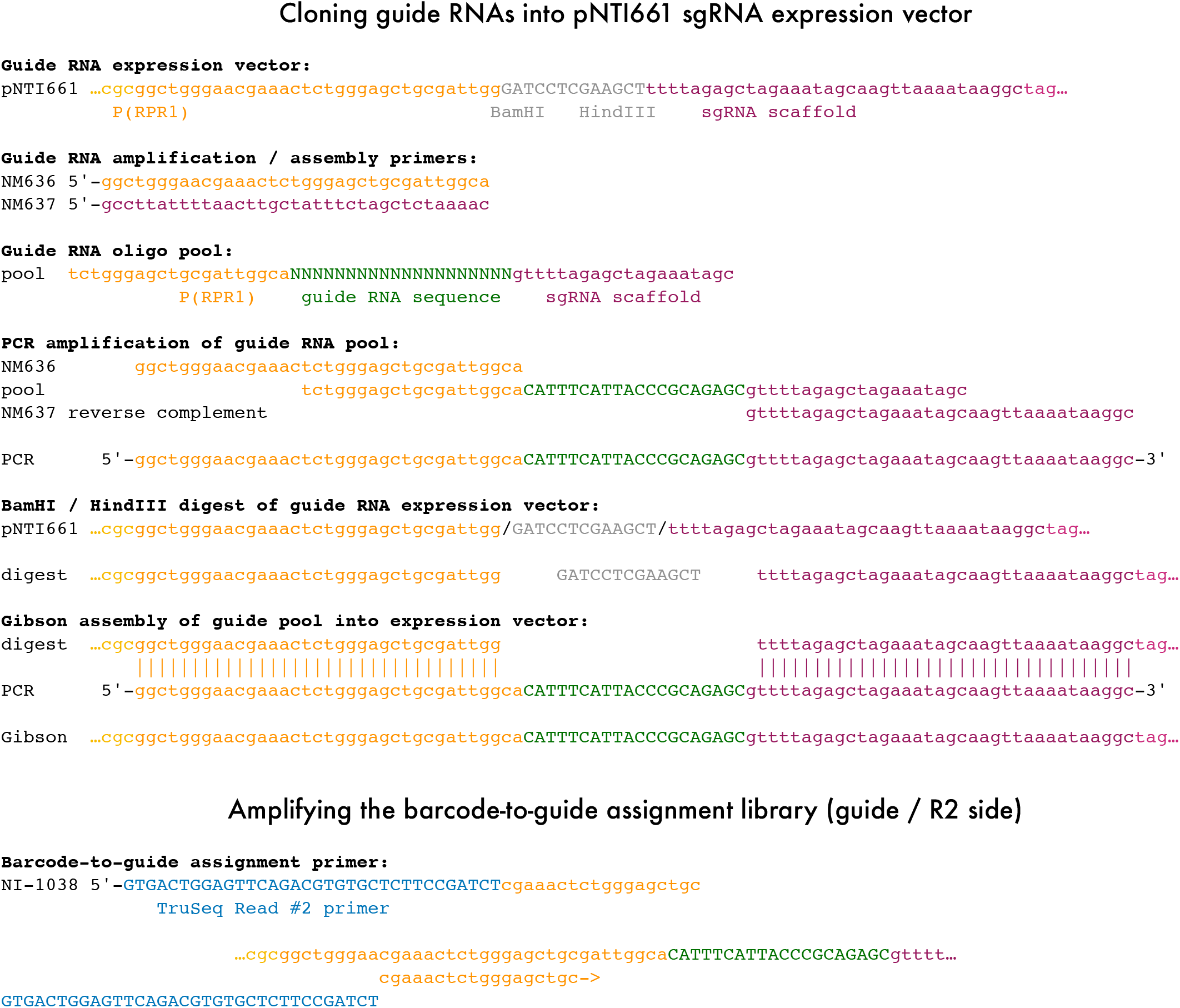
Guide RNA cloning strategy. The guide RNA expression vector is linearized by digestion with BamHI and HindIII. The guide RNA oligonucleotide pool is amplified by PCR with NM636 and NM637 primers to create a substrate for Gibson assembly. The assembly reaction reconstitutes an intact *P(RPR1)-sgRNA* expression cassette. PCR using NI-1038 will prime in the *P(RPR1)* region and amplify a fragment that includes the variable guide RNA sequence flanked by a portion of one Illumina sequencing adapter, corresponding to the P7 side of the library with the Read #2 primer site.

**Figure S2.**
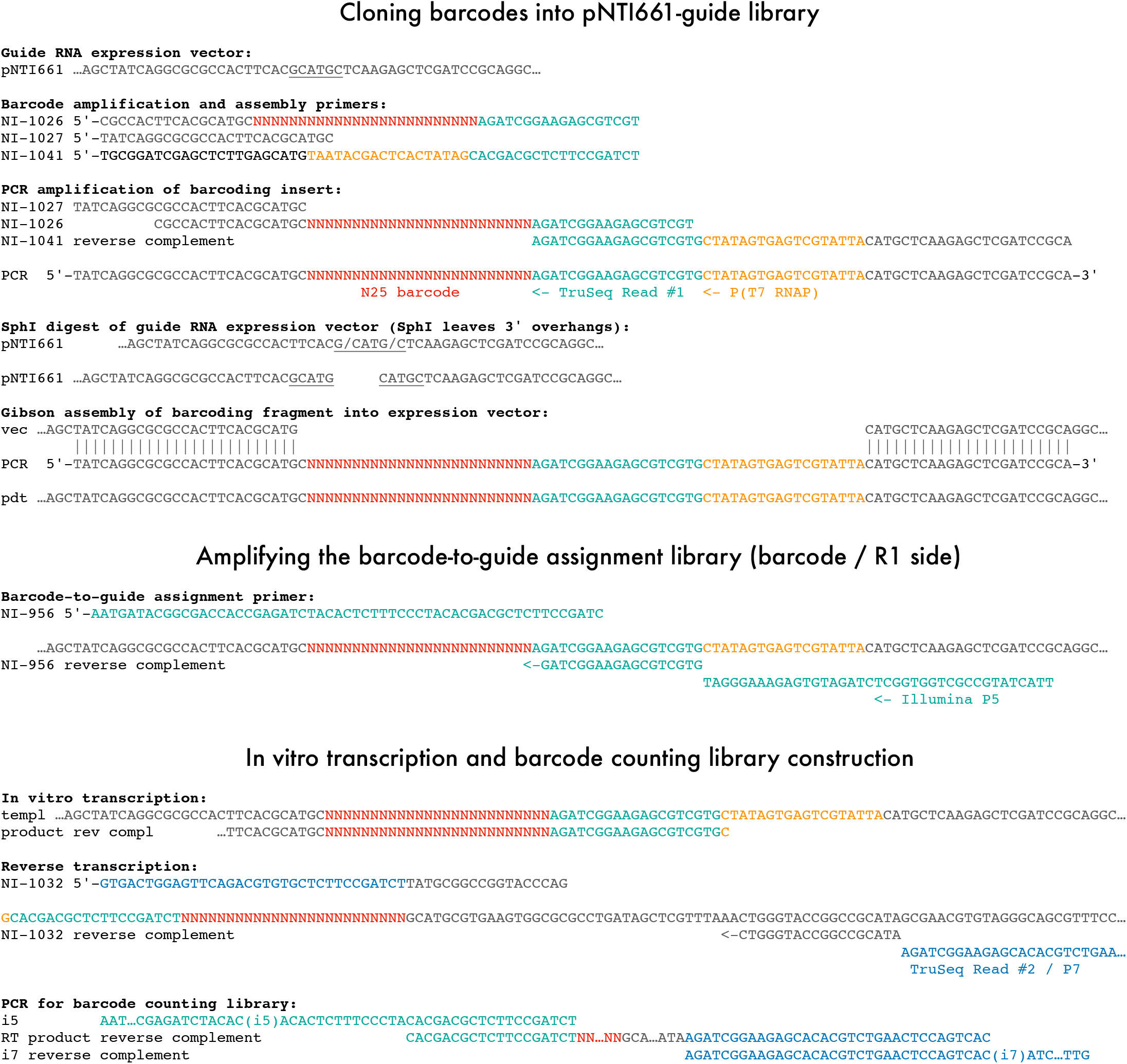
Barcode cloning strategy. The guide RNA expression vector is linearized by digestion with SphI, an enzyme that leaves 3’ overhangs that are not resected during Gibson assembly. The barcode insert is amplified by PCR from one oligonucleotide, NI-1026, containing a random N_25_ sequence, with flanking primers NI-1027 and NI-1041 that add flanking regions suitable for cloning. After assembly into the SphI site, the barcode is flanked immediately by a fragment of the Illumina TruSeq Read #1 site, with a T7 RNA polymerase promoter located just beyond this primer site. PCR using NI-956 will prime on this TruSeq Read #1 sequence and amplify a fragment that includes the barcode flanked by an Illumina sequencing adapter, corresponding to the P5 side of the library. In vitro transcription produces an RNA that includes the Read #1 site. Reverse transcription with a specific primer containing the TruSeq Read #2 / P7 sequence yields a cDNA that can be amplified into an Illumina library that includes the barcode.

**Figure S3.**
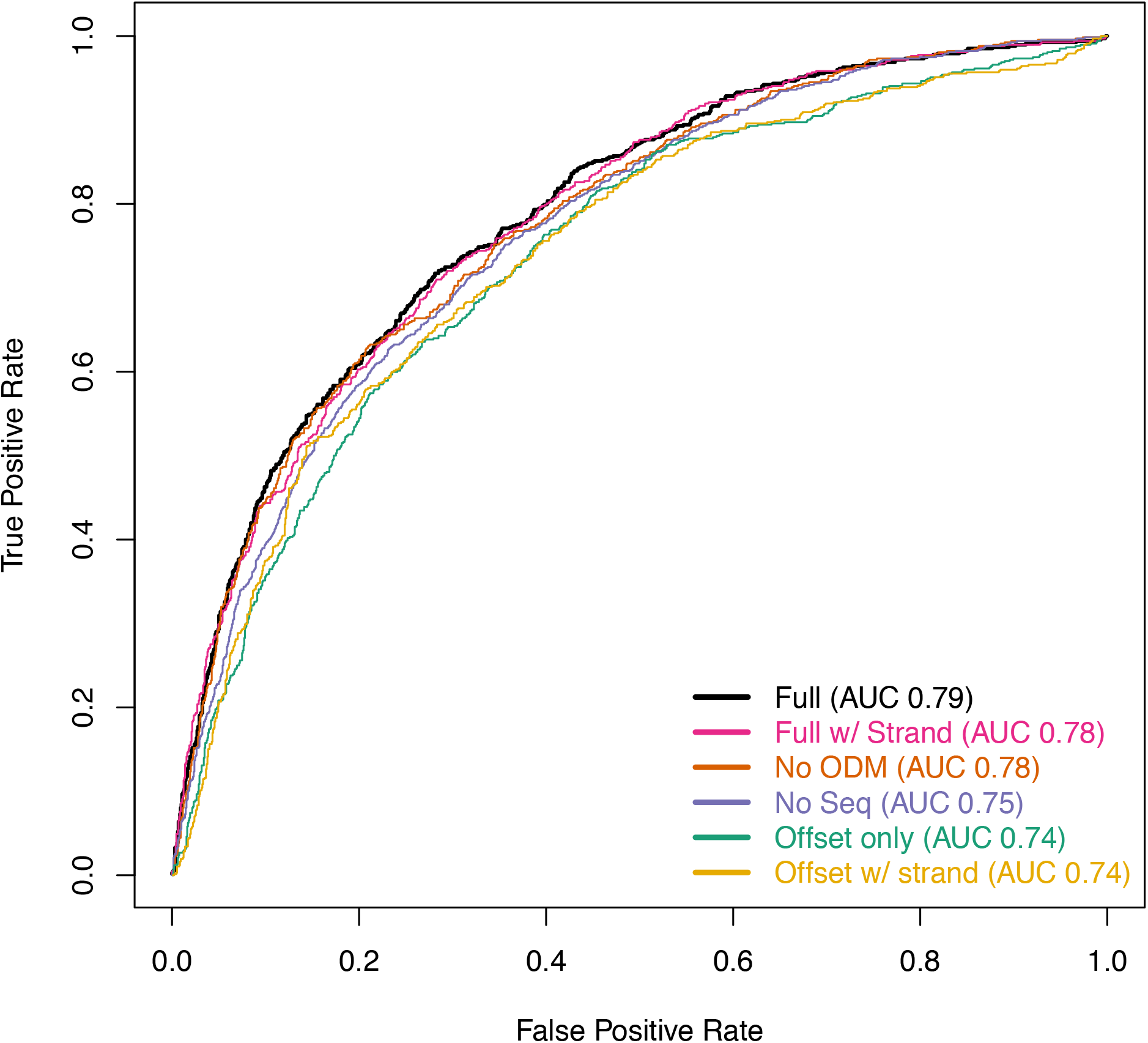
Accurate predictions of guide activity. Receiver operating characteristic for logistic regression models of guide activity. The full model includes methylation-based accessibility data (ODM-Seq) and guide sequence along with the position of the guide relative to the transcription start site. The “No ODM” and “No Seq” models exclude ODM-Seq and guide sequence parameters, respectively, and “Offset only” excludes both. The models “w/ Strand” include separate local regressions for guide activity based on the strand of the guide site relative to the target gene. All AUC values are calculated in the same 10-fold cross validation analysis.

**Table S1.**
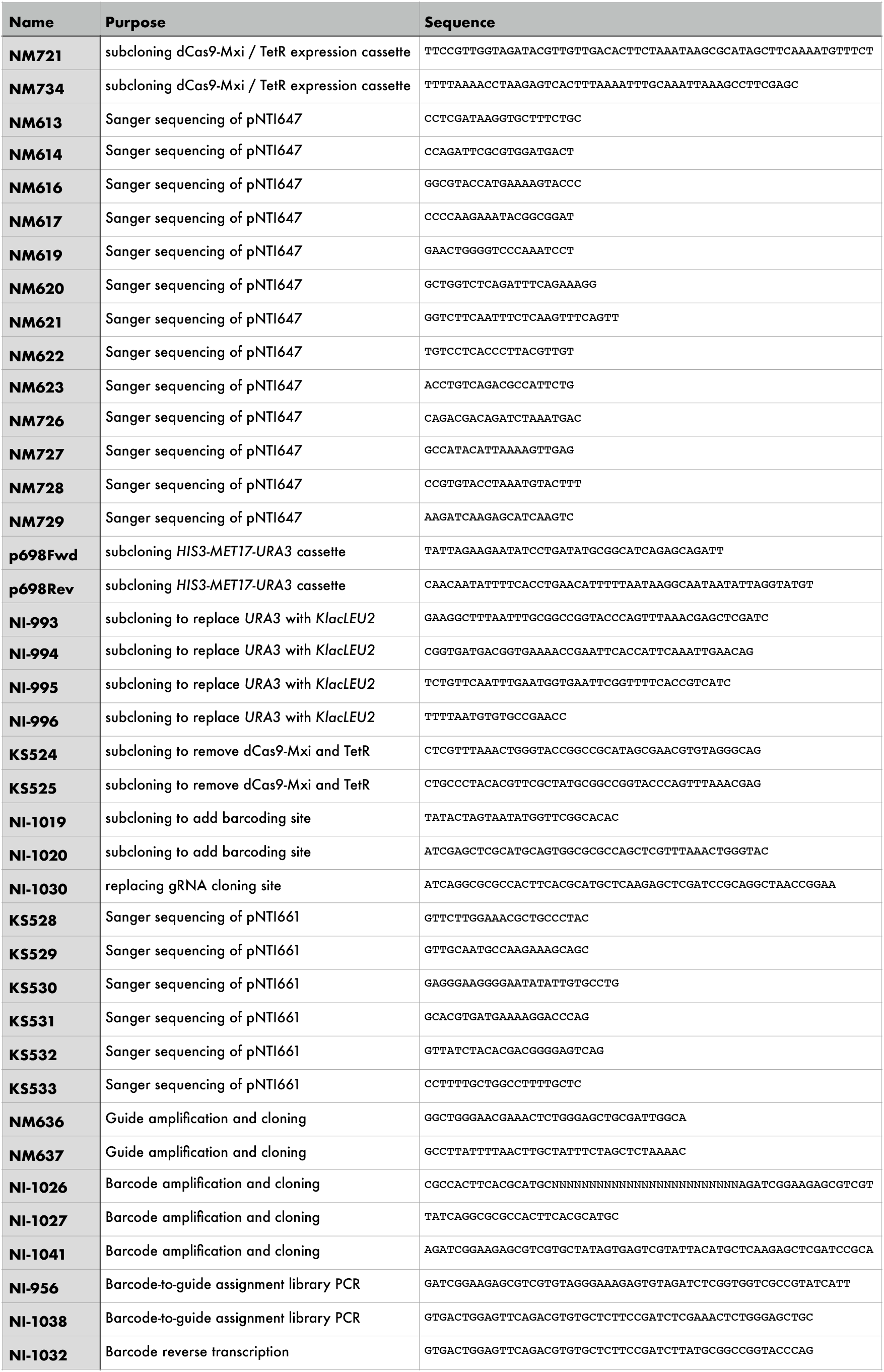
Oligonucleotide sequences used in this study.

